# SARS-CoV-2 detection from nasopharyngeal swab samples without RNA extraction

**DOI:** 10.1101/2020.03.28.013508

**Authors:** Carolina Beltrán-Pavez, Chantal L. Márquez, Gabriela Muñoz, Fernando Valiente-Echeverría, Aldo Gaggero, Ricardo Soto-Rifo, Gonzalo P. Barriga

## Abstract

The ongoing COVID-19 pandemic has reached more than 200 countries and territories worldwide. Given the large requirement of SARS-CoV-2 diagnosis and considering that RNA extraction kits are in short supply, we investigated whether two commercial RT-qPCR kits were compatible with direct SARS-CoV-2 detection from nasopharyngeal swab samples. We show that one of the tested kits is fully compatible with direct SARS-CoV-2 detection suggesting that omission of an RNA extraction step should be considered in SARS-CoV-2 diagnosis.

## Overview

Two coronaviruses, SARS-CoV (severe acute respiratory syndrome coronavirus) and MERS-CoV (Middle East respiratory syndrome coronavirus) caused large-scale severe respiratory syndromes in humans in 2002 and 2013, respectively ^1, 2^. A recent outbreak of pneumonia in Wuhan, China declared in December 12^th^2019 led to the isolation and identification of a new human coronavirus, 2019-nCoV (SARS-CoV-2), as the responsible agent of this new acute respiratory syndrome ^3^. Since, SARS-CoV-2 and the Coronavirus disease 19 (COVID-19) have spread to more than 200 countries and territories worldwide affecting more than 600,000 people (laboratory-confirmed cases) from which around 28,000 have died by COVID-19 ^4^. As such, the WHO declared COVID-19 outbreak as a pandemic on March 11^th^2020 ^5^. Given the rapid spread of the SARS-CoV-2 virus and that an important fraction of the infected individuals remains asymptomatic but still spread the virus ^6^, rapid and extensive diagnosis together with the isolation of infected individuals remains as one of the most powerful ways to avoid the spread of new COVID-19 cases. There are several RT-qPCR protocols from different countries that are currently approved by the WHO in order to detect SARS-CoV-2 ^7^. However, one of the limiting steps of such protocols is RNA extraction from clinical samples, which is currently performed by the use of a limited quantity of approved RNA extraction kits. Since the current demand for those kits largely exceeds their offer, it is necessary to develop alternative strategies allowing a rapid detection of the virus. This becomes more important in middle- and low-income countries, which will not have massive access to kit-based diagnosis ^8, 9^. In this sense, it was recently reported that the detection of SARS-CoV-2 by RT-qPCR from a mixed sample could be achieved with non-approved kits but also in the absence of an RNA extraction process ^10^. Here, we have extended these preliminary observations by using an RNA extraction-free SARS-CoV-2 detection protocol from nasopharyngeal swab samples from two COVID-19 diagnosed individuals using two commercially available and broadly used RT-qPCR kits. Our data indicate that RNA extraction can be omitted from the protocol at least with one of the tested kits allowing the rapid and reliable detection of SARS-CoV-2 directly from nasopharyngeal swabs samples.

## Comparison of SARS-CoV-2 detection using RNA or direct nasopharyngeal swabs samples

Nasopharyngeal swabs samples (NSS) from two laboratory-diagnosed COVID-19 positive individuals (both confirmed by the Chilean Institute of Public Health) were obtained from the Servicio de Laboratorio Clínico, Hospital Clínico de la Universidad de Chile “Dr. José Joaquín Aguirre”, Santiago, Chile. FLOQswabs (Copan Diagnostics Inc) containing the nasopharyngeal samples were added to a 4 ml tube containing 3 ml of UTM-RT mini transport media (Copan Diagnostics Inc). A volume of 5 μl of the NSS was used to perform the RT-qPCR detection using the TaqMan™ 2019-nCoV Assay Kit v1 (ThermoFisher) and the 2019-nCoV CDC qPCR Probe Assay (Integrated DNA Technologies). RT-qPCR detections using 5 μl of RNA extracted with the QIAamp Viral RNA Mini Kit (Qiagen) were processed in parallel in order to perform comparisons. In both cases, the cycling protocol recommended by each supplier was used in the QuantStudio3 Real Time PCR System (Thermo Fisher Scientific) using the samples obtained from the two COVID-19 positive individuals, the positive control provided by each kit together with SARS-CoV-2-free RNA obtained from human embryonic kidney (HEK293T) cells, which was used as a negative control.

We first tested the suitability of the TaqMan™ 2019-nCoV Assay Kit v1 to detect the SARS-CoV-2 genes Orf1ab, S and N (Fig. 1A and Table 1). We observed that the three viral genes were readily detected in the NSS of both COVID-19 positive samples and the positive control but not in RNA from HEK293T cells. Interestingly, detection of SARS-CoV-2 genes from the NSS was as efficient as RNA samples extracted with the QIAamp Viral RNA Mini Kit with differences between 1 to 5 Ct values depending on the viral gene and the sample (Fig. 1A and Table 1). These results strongly indicate that the TaqMan™ 2019-nCoV Assay Kit v1 from Thermo Fisher Scientific is fully compatible with the direct use of NSS without any RNA extraction step.

**Table 1.** Comparative Ct values in kit-extracted RNA and direct nasopharyngeal swab samples (NSS).

|  | Patient 1 |  | Patient 2 |  | HEK 293 cells |
| --- | --- | --- | --- | --- | --- |
|  | RNA | NSS | RNA | NSS | RNA |
| **TaqMan 2019-nCoV** |  |  |  |  |  |
| Orf1ab | 26,346 | 27,045 | 18,148 | 19,322 | Undetermined |
| RNaseP | 25,889 | 30,196 | 20,415 | 27,619 | 9,142 |
| N | 25,602 | 26,813 | 17,408 | 19,153 | Undetermined |
| RNaseP | 25,727 | 30,561 | 20,133 | 27,332 | 8,749 |
| S | 24,210 | 27,510 | 15,716 | 20,49 | Undetermined |
| RNaseP | 25,788 | 30,695 | 19,989 | 27,216 | 10,136 |
| **2019-nCoV CDC** |  |  |  |  |  |
| N1 | 24,260 | 28,398 | 15,547 | 21,654 | 38,124 |
| N2 | 25,904 | 38,121 | 17,648 | 30,268 | Undetermined |
| RNaseP | 29,761 | 31,166 | 25,917 | 29,55 | 20,582 |

**Figure 1:**
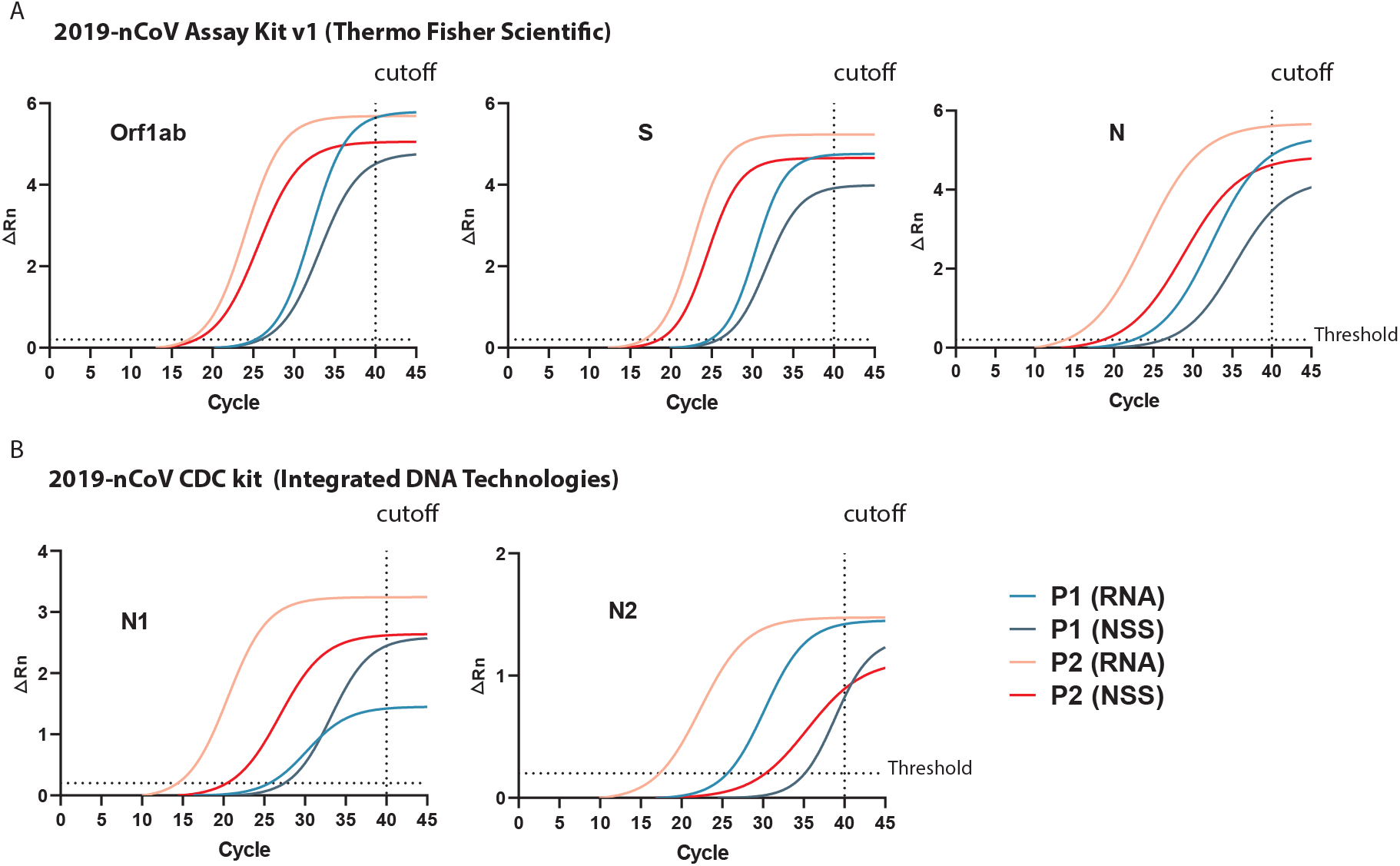
Detection of SARS-CoV-2 from nasopharyngeal swab samples (NSS) or kit-extracted RNA (RNA) from two COVID-19 laboratory-diagnosed patients (P1 and P2). RT-qPCR reactions were carried with the 2019-nCoV Assay Kit v1 from Thermo Fisher Scientific (A) or the 2019-nCoV CDC kit (Integrated DNA Technologies) in a QuantStudio3 Real Time PCR System (Thermo Fisher Scientific) following suppliers’ indications for cycling. Data were analyzed with the QuantStudio3 Real Time PCR Software and Fitted line plotted using a non-linear regression fit model with the GraphPad Prism 8.1 software. Dotted lines indicate the threshold of 0,2.

We then wanted to know whether these observations were extrapolable to the 2019-nCoV CDC qPCR Probe Assay (Fig. 1B and Table 1). As observed, although we were able to detect the SARS-CoV-2 N gene from the NSS using the N1 and N2 probes provided by the kit, our data indicate that this kit is much more performant using RNA extracted samples. This observation was more evident with the N2 probe, in which we observed a difference of 13 Ct values between the NSS and the RNA extracted samples from both patients (Fig. 1B and Table 1).

## Conclusions

In summary, we have assayed the suitability of the TaqMan™ 2019-nCoV Assay Kit v1 from Thermo Fisher Scientific and the 2019-nCoV CDC qPCR Probe Assay from Integrated DNA Technologies to directly detect SARS-CoV-2 RNA from nasopharyngeal swabs samples from two COVID-19 positive individuals diluted in transport media. Our results indicate that the kit from Thermo Fisher Scientific is fully compatible with an RNA extraction-free protocol allowing the detection of viral RNA with an efficiency comparable to that obtained using kit-extracted RNA. Our observations will be helpful to support SARS-CoV-2 diagnoses in places such as the USA and Western Europe where COVID-19 cases have exploded during the last weeks but also in middle- and low-income countries, which would not have a massive access to kit-based diagnosis.

## Ethical statement

Given the important impact that data from this study could have on Public Health, the two nasopharyngeal samples from COVID-19 positive patients were deidentified and not considered as Human samples. We are currently, preparing a protocol to be presented at the Ethic Committee of the Faculty of Medicine at Universidad de Chile in order to improve SARS-CoV-2 detection strategies using a larger number of Human samples.

## Acknowledgements

The authors are supported by ANID Chile through Fondecyt grants Nº 1181656 (AG), 1190156 (RS-R), 1180798 (FV-E), 3200556 (CLM); Instituto Nacional Antártico de Chile RT_35-19 (GB-P); Proyecto de Internacionalización UCH-1566 (CB-P). Authors wish also thanks to Thermo Fisher Scientific Chile for providing technical support and the Chilean Science, Technology, Knowledge and Innovation Ministry for articulating and coordinating support from the scientific community.

## Authors contributions

CB-P, CLM, FV-E, AG, RS-R and GB participated in the study design. CB-P, CLM and GB performed the experiments. CB-P, CLM, RS-R and GB analysed the data. GM provided the NSS from COVID-19 diagnosed individuals. FV-E, AG, RS-R and GB wrote the manuscript. All authors approved the final version of the manuscript.

## Conflict of interest

The authors declare that there are no conflicts of interest associated with this work.

